# Agonistic interactions between slender mudskippers (*Periophthalmus gracilis*) in Singapore

**DOI:** 10.1101/2025.01.27.635183

**Authors:** Soroush Saleh, Tan Alyssa Yan Ying, Edmund Chow Wei Jie, Chee Koi Jun, Sean Neville, Philip Johns

## Abstract

Mudskippers engage in frequent territorial interactions. Few studies of mudskipper agonistic behavioral sequences exist. The slender mudskipper (*Periophthalmus gracilis*) inhabits intertidal zones of mangrove forests across Southeast Asia. Here, we describe common behavioral sequences of winners and losers among unmanipulated, wild slender mudskipper interactions in Singapore. Winning mudskippers showed more approach behaviors than losers, such as “Crawl towards”, and more behaviors that may indicate aggression, such as “Pectoral wave”, “Flag”, and “Head up”. Losers showed more evasive behaviors, like “Jump” and “Crawl away”. We found evidence of agonistic behavioral matching. Larger mudskippers won contests more frequently than smaller mudskippers.

## Introduction

Territorial disputes and aggressive interactions constitute a sizeable proportion of animal social interactions in some animals (Peake and McGregor, 2004). Aggressive interactions can surface in social environments where competitors stand to gain resources such as food, shelter, territory, and access to mates (Arnott and Elwood, 2008; Hinsch and Komdeur, 2017; Peake and McGregor, 2004), and enable each competitor to assess each other’s fighting ability (Arnott and Elwood, 2009; Peake and McGregor, 2004; Reddon et al., 2013). Fishes may display different contest behaviors depending on their assessment of a contested resource’s value and their internal state (Hsu et. al., 2011).

Mudskippers are intertidal fish that forage for insects, worms, and small crustaceans at low tides in mangroves and mudflat ecosystems (Dinh et al., 2021; Hui et al., 2019). Prior studies of mudskipper social behaviors describe territorial displays and agonistic encounters (e.g., Brillet 1983, Clayton and Townsend 2017; Clayton and Vaughan 1988; Townsend and Tibbetts, 2005) but not detailed systematic analyses of agonistic behavioral sequences. Slender mudskippers, *Periophthalmus gracilis*, are common in the coastal mudflats in Southeast Asia, including in Singapore, Indonesia, Malaysia, Thailand, Vietnam and Philippines (Dinh et al., 2021; Larson et al., 2008; Murdy, 1989; Taniwel et al., 2020).

Here we describe agonistic interactions in the slender mudskipper, *P. gracilis*, in Singapore. We include behaviors not previously described in this species (see Clayton and Townsend, 2017). Our aim is to identify influences on winning and losing individuals, including differences in behavioral frequencies and behavioral sequences between winners and losers. Previous research on five sympatric species in Australia, which included the *P. gracilis*, suggest that mudskipper size and dominance hierarchy may determine interaction outcomes (Nursall, 1981). Our research examines whether relative size is a determinant of the outcomes of agonistic interactions between *P. gracilis*.

## Materials and Methods

We videorecorded mudskipper interactions at Berlayer Creek in Labrador Nature Park in Singapore (1.271705ºN, 103.802992ºE to 1.266566ºN, 103.806168ºE), at low tides during daylight hours when the mudflats were exposed, from 11 February to 14 March 2023. Slender mudskippers are about 4.4 cm long (Hua et al. 2023) and are recognizable by the irregular blackish dots on the first dorsal fin (Hua et al., 2023; Jaafar et al., 2016; Murdy, 1989), with several silvery white vertical lines on the trunk stretching from the pectoral-fin base to the caudal peduncle (Jaafar et al., 2016). At three observation platforms along Berlayer Creek, we scanned the mudflat on the opposite shore 5-10m away for presence of mudskippers. Once we spotted a mudskipper, we recorded all behaviors for 1 minute using consumer-grade cameras (Nikon p900 Coolpix (https://www.nikon.com) and Canon EOS 90D (https://global.cnon/en/). If the mudskipper engaged in aggressive interactions, we continued recording all behaviors of both mudskippers until the end of the encounter. Data are archived at DOI: 10.6084/m9.figshare.24781932.

We defined the start of an interaction as the point when one individual changed its orientation or performed a behavior in direct response to another individual’s movement. We defined the end of an interaction as the point at which one mudskipper left the site of conflict without the other mudskipper pursuing it, or when one or both mudskippers turned away from each other and ceased movement. We defined losers of interactions as the mudskipper that moved the farthest away from the initial site of the interaction and defined the winner as the mudskipper that remained closest to the site. We identified mudskippers that did not respond to the behaviors of other mudskippers in the video frame as “neutrals.”

We identified repeatable behaviors (see Table S1, Fig. S1). All behaviors except “Contact”, “Crawl towards”, “Crawl away”, “Wiggle”, and “Blink” (Table S1) were taken from Clayton and Townsend’s (2017) ethogram. We scored videos of interactions between winners and losers using BORIS (Friard and Gamba, 2016; www.boris.unito.it). We described behavioral sequences with the BORIS program Behatrix. (www.boris.unito.it/pages/behatrix), using 1000 permutations to determine which transitions occurred more frequently than chance. We visualized behavioral transitions in kinetic flow diagrams using GraphViz (Ellson et al., 2002; https://graphviz.org).

We used R for statistical analyses (R Core Team, 2023; https://www.R-project.org/). We compared the frequencies of behaviors, in aggregate, of the winners and losers with a permutated multiple analysis of variance (PERMANOVA), using the function *adnois2* (Anderson, 2017; R vegan package, https://github.com/vegandevs/vegan) and 1000 permutations. To account for the pairwise nature of interactions, we nested the behavioral frequencies of winners and losers within individual encounters with the *strata* argument. In this study, we used a distance matrix (*dismatrix*) because the research question focused on pairwise dissimilarities (Anderson, 2017).

We compared relative sizes of winners and losers within each interaction from video images using ImageJ (Schneider et al. 2012; https://imagej.net/ij/). Because our sampling procedure was non-invasive, we did not directly measure Total Length (**TL**) of the mudskipper (Fig S2). For each mudskipper, we took screen captures of the videos, averaged three measurements of TL, and compared losers’ TL as percentages of the winners’ TL. We could not ascertain sex from video observations and did not include that information in this this study.

## Results

We collected data on 49 complete interactions. The duration of interactions averaged 10.17s (± 6.97s SD). Behavioral analysis revealed different behavioral sequences for winning and losing mudskippers (Fig. 1). Winners (Fig 1a) repeated Pectoral wave behaviors (77.8%). Winners always performed Head up before Wiggle (100%). Winners sometimes performed Flag, an agonistic behavior (Clayton and Vaughan, 1988), before Contact (25%). This suggests that a Flag signaled an escalation in the encounter. Blink always led to Flip (100%). Flip sometimes led to Crawl away (15.8%), and often led to Jump (63.2%). Winners performed Crawl away repeatedly (28.6%) less often than losers did (57.8%), which may be a result of the winners returning from the periphery to the center of a territory after a fight ended, resulting in winners and losers retreating from the same area.

**Figure 1.**
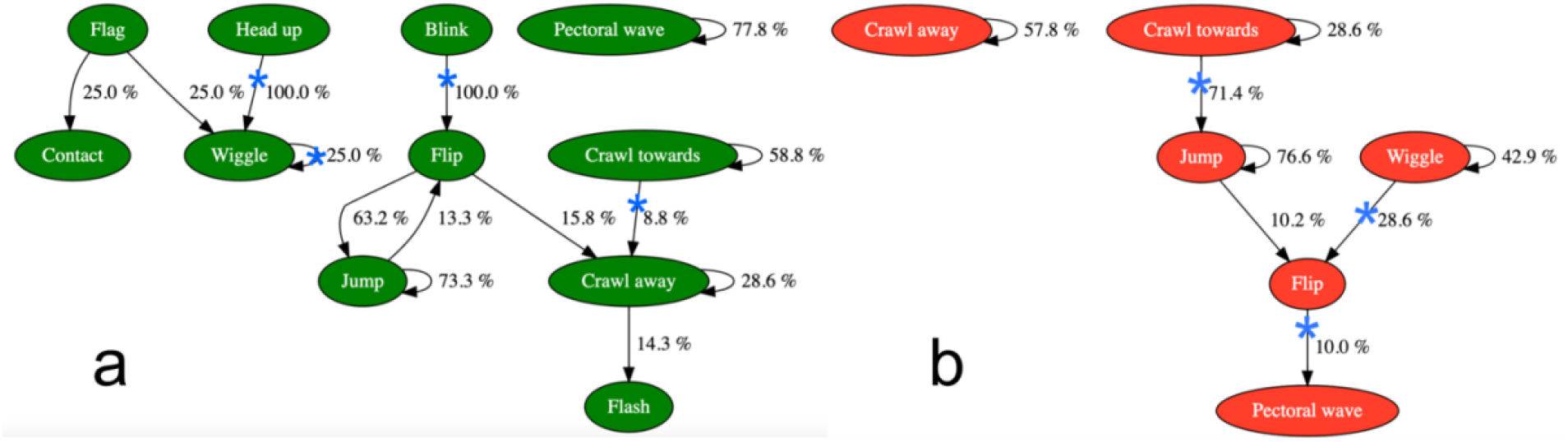
Behavioral transitions for winners (green; a), losers (red; b) that occurred more frequently than chance as determined by permutation test (see text). Blue asterisks denote marginal significance (0.05 < p < 0.06).

Among losers (Fig 1b), Crawl away often led to more Crawling away (57.8%), which suggests retreat. Crawl towards sometimes led to Crawl towards (28.6%), as fish moved closer towards each other, suggesting escalation by the loser. Also, Jump often followed Crawl towards (71.4%), which might signal the beginnings of agonistic behavior by the eventual losers. Losers often Jumped repeatedly (76.6%). They also Wiggled repeatedly (42.9%), and sometimes followed a Wiggle with a Flip (28.6%). Lastly, the Flip sometimes led to a Pectoral wave (10.0%).

We described the behavioral sequences of winning and losing mudskippers together (Fig. 2). The loser’s Flag display always preceded the winner’s Flag (100%). The loser’s Flip occasionally led to the winner’s Wiggle (11.1%). The loser’s Crawl away led to the same action repeated almost half the time (43.6%), suggesting a retreat. The winner’s Jump very rarely preceded the winner’s Quiver (1.5%), but it often led to the loser’s Jump (50.4%).

**Figure 2.**
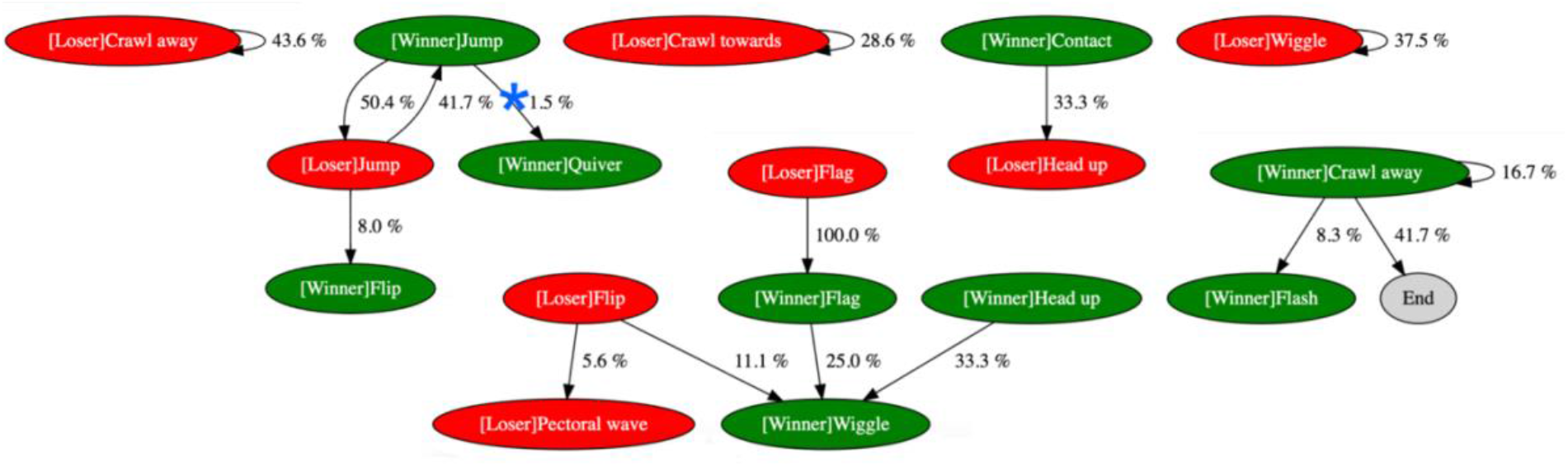
Behavioral transitions for winners (green) and losers (red), combined, that occurred more frequently than chance, as determined by permutation test (see text). Blue asterisks denote marginal significance (0.05 < p < 0.06).

Because mudskippers respond to the behaviors of competing individuals, we compared the differences in the frequencies of behaviors between winners and losers paired within interactions. Winners and losers within interactions performed behaviors with different frequencies, in aggregate (PERMANOVA F_1,96_ = 5.10, p < 0.001), although the differences accounted for relatively little of the overall variation in behavioral frequency (R^2^ = 0.050). Losing mudskippers performed more frequent Crawl away and Jump behaviors than winners; winning mudskippers exhibited more frequent Head up and Flag behaviors than losers (Fig. 3).

**Figure 3.**
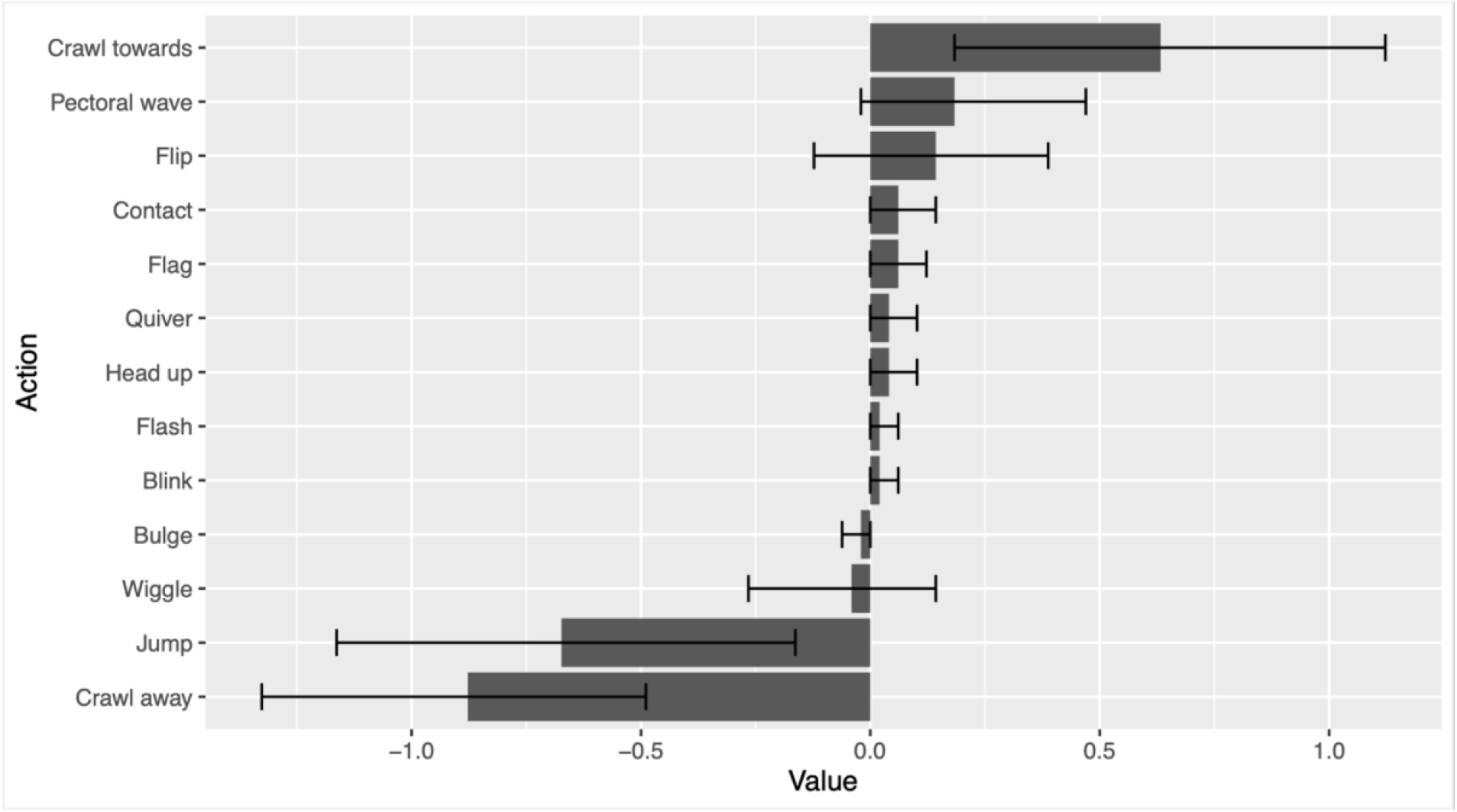
Average paired differences between winner and loser behavioral frequencies within interactions. Negative values indicated losers performed a behavior more frequently; positives values indicate winners performed a behavior more frequently. Error bars indicate bootstrapped estimates (r = 1000) of 95% confidence intervals.

On average, losing slender mudskippers performed more total behaviors (5.24 ± 0.42 SE) than did winning mudskippers (4.80 ± 0.31 SE), although there was no significant pairwise differences in the overall number of behaviors performed by winners and losers (Wilcoxon signed rank test, V = 379, p = 0.256. Larger mudskippers (n = 44) won contests significantly more frequently than smaller mudskippers (chi-squared = 63.38, df = 1; p < 0.001). Regression analysis revealed no significant effect of the percent size difference between winners and losers on the duration (in seconds) of aggressive interactions (F_1,47_ = 0.918, p = 0.344,).

## Discussion

Some components were similar between winning and losing mudskippers. Both winners and losers perform the sequence, Jump → Flip. In general, the losers’ actions are locomotory (crawling towards or away), leading to more crawling towards or away (see Fig. 1b). This finding is consistent with our observations of losing mudskippers travelling through the territories of winning mudskippers, and often being chased off. Successive Crawl away behaviors by the losing mudskipper occurred about twice as often as by the winners and may be an evasive behavior that can resolve the encounter. It is interesting that winners sometimes performed Flash after the Crawl away (14.3%), since Flash may be an agonistic behavior (Clayton and Vaughan, 1988). Among winners, Crawl towards led to repeated Crawl towards about twice as frequently (58.8%) as among losers (28.6%), which is consistent with winners continuing to escalate agonistic encounters. Winners followed Head up with Wiggle, and followed Blink with Flip, every time. It isn’t clear whether the sequential nature of these behaviors is communicative, or if there is a biomechanical or physiological reason. In contrast, winning mudskippers often perform what may be threatening behavioral sequences, such as Wiggle → Pectoral wave (Fig. 1a), which could function to advertise or exaggerate mudskipper size.

The winners Flag display always followed the losers Flag (100%), which suggests behavioral matching. This observation in slender mudskippers is consistent with agonistic behavior in *B. dussumieri* (Clayton and Vaughan, 1988), territorial behavior of *B. boddarti* (Swennen et al., 1995) and aggression of *S. histophorus* (Aguilar, 2000). This sequence suggests that slender mudskippers assess each other in contests.

Pairwise comparisons reveal that winners and losers perform behaviors with different frequencies in aggregate, although those differences accounted for a small percentage of the variation. The magnitude of the differences, in aggregate, might suggest that behavioral sequences contribute more to interaction outcomes than behavioral frequencies, *per se*, although obviously the two are not independent. Winning mudskippers performed some individual behaviors more frequently than losing mudskippers, such as Crawl towards, Pectoral wave, Contact, and Flag (Fig. 3), and these behaviors may function to or escalate interactions. Notably, winners followed losers Flag displays with Flags of their own, but they also performed Flag behaviors more than losers, and that led to contact (Fig 1a), which suggests escalation. Losers Jump and Crawl away more than winners. The tendency for losers to crawl away is partly due to our definition of the interaction loser. Pairwise comparisons also raise questions about the function of Jump, *per se*. In both winners and losers, Jump often preceded the end of an interaction. Whether the Jump behavior is assessment or escalation is not clear from our analysis.

Because our study was non-invasive, we only measured the relative size of mudskippers in aggressive interactions. Larger mudskippers almost always won interactions, which suggests that resource-holding potential is related to size. However, we found no significant relationship between percent size difference and the duration of interactions, i.e., much larger mudskippers did not resolve fights more quickly than marginally larger mudskippers. There might be other contributors to outcome and interaction duration, including the sex and age of the mudskippers. In some species of mudskippers, aggressive behaviors like raising fins can also serve as a courtship signal (Jaafar and Murdy, 2017). While there are external differences between male and female slender mudskippers in the color of the first dorsal fin (Jaafar and Murdy, 2017), we were generally unable to tell males from females *in situ* through camera lenses from several meters away. The first dorsal fin was rarely raised, and even when it was, it was hard to assess coloration of muddy mudskippers in natural light.

Our research constitutes the first detailed analysis of the agonistic behavioral sequences of *P. gracilis*. Despite limitations, we observed a strong relationship between size and agonistic outcome (win or lose) of *P. gracilis*. This result concurs with previous research conducted by Brillet (1981) who noted that larger *P. argentilineatus* were mostly victorious (94%-100%) and initiated more agonistic encounters. Fin raising has been identified as an important behavior in agonistic interactions of mudskippers, being observed more frequently in subordinate mudskippers (Jaafar and Murdy, 2017). Laboratory experiments involving the removal of dorsal fins in *P. argentilineatus* has resulted in dominance (Brillet, 1984) suggesting that the main agonistic purpose of this behavior is to signal submissiveness. The tendency of losers to raise both fins after contact and winners to subsequently skip away suggests that fin raising in *P. gracilis* may act as a surrender signal. Further studies are required to better understand the specific nature of fin raising in agonistic interactions of mudskippers.

## Acknowledgements

Portions of this study were conducted by Saleh, Tan, Chow, Chee, and Neville, in partial fulfilment of the requirements for the Urban Wildlife Studies, and the Independent Reading and Research, courses at Yale-NUS College. The work was partly supported by the Singapore Ministry of Education through the Yale-NUS College grant R-607-265-226-121.

## Data availability

Data are archived at DOI: 10.6084/m9.figshare.24781932.

## Ethical approval

This study was purely observational. Animals were not captured, handled, manipulated, or fed in any way. All observations occurred several meters away from subjects. The study therefore required no Yale-NUS College IACUC approval.

