## Supplementary material for "Agonistic interactions between slender mudskippers (*Periophthalmus gracilis*) in Singapore": Table S1

**Supplementary Table**

Table S1. Ethogram describing behaviors[a] displayed by *P. gracilis* in paired fights.

| No. | Behaviour | Description |
|---|---|---|
| 1 | Jump | Skip |
| 2 | Flip | Lateral flip |
| 3 | Crawl towards | Crawl towards other mudskipper |
| 4 | Crawl away | Crawl away from other mudskipper |
| 5 | Head up | Head raised |
| 6 | Flag | First dorsal fin raised |
| 7 | Flick | Second dorsal fin raised |
| 8 | Flash | Both dorsal fins raised |
| 9 | Pectoral wave | Movement in pectoral fin |
| 10 | Quiver | Body waving |
| 11 | Contact | Physical contact with other mudskipper |
| 12 | Wiggle | Sideways movement while remaining stationary |
| 13 | Bulge | Cheeks puffed |
| 14 | Blink | Brief closing and opening of eyes |

| | | |
|---|---|---|
| 15 | Roll | Dropping on its flank |

a. Clayton & Townsend (2017) originally described “Jump”, “Flip”. “Head up”, “Flag”, “Flick”, “Flash”, “Pectoral Wave”, “Quiver” and “Bulge”; remaining behaviors are newly added.
