## Supplementary material for "Agonistic interactions between slender mudskippers (*Periophthalmus gracilis*) in Singapore": Fig. S1

**Supplementary Figure**

Fig. S1. Behaviors recorded during study as additional behaviors to cited ethogram (Table S1). Panels displayed from left to right depict the progression of behavior **a** Contact recorded between two individuals in second panel and end of contact is recorded in third panel. **b** Wiggle recorded in an individual.

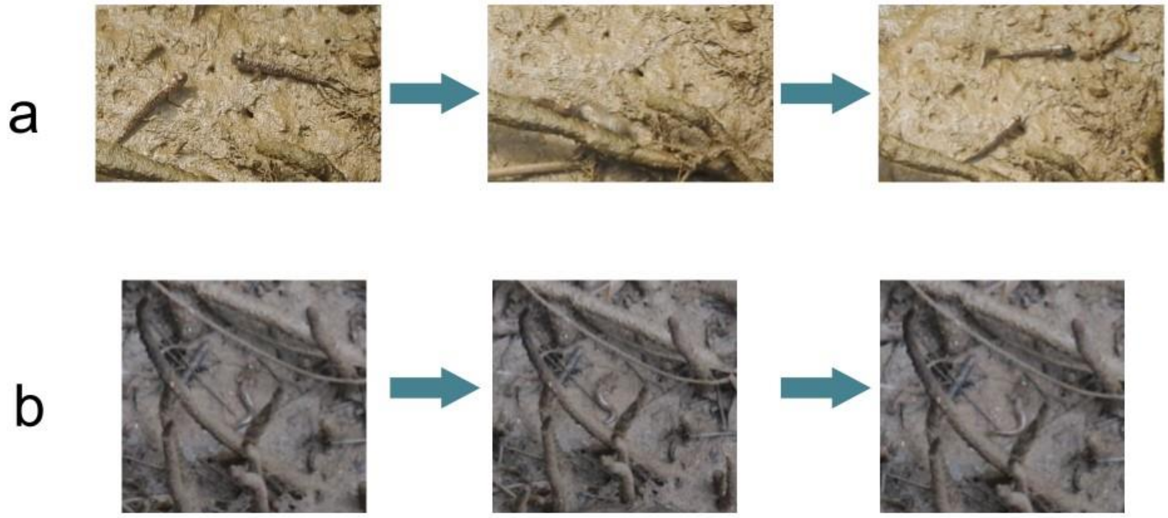
